# Recurrent Evolutionary Innovations in Rodent and Primate *Schlafen* Genes

**DOI:** 10.1101/2024.01.12.575368

**Authors:** Joris Mordier, Marine Fraisse, Michel Cohen-Tannoudji, Antoine Molaro

## Abstract

SCHLAFEN proteins are a large family of RNase-related enzymes carrying essential immune and developmental functions. Despite these important roles, *Schlafen* genes display varying degrees of evolutionary conservation in mammals. While this appears to influence their molecular activities, a detailed understanding of these evolutionary innovations is still lacking. Here, we used in depth phylogenomic approaches to characterize the evolutionary trajectories and selective forces shaping mammalian *Schlafen* genes. We traced lineage-specific *Schlafen* amplifications and found that recent duplicates evolved under distinct selective forces, supporting repeated sub-functionalization cycles. Codon-level natural selection analyses in primates and rodents, identified recurrent positive selection over Schlafen protein domains engaged in viral interactions. Combining crystal structures with machine learning predictions, we discovered a novel class of rapidly evolving residues enriched at the contact interface of SCHLAFEN protein dimers. Our results suggest that inter Schlafen compatibilities are under strong selective pressures and are likely to impact their molecular functions. We posit that cycles of genetic conflicts with pathogens and between paralogs drove Schlafens’ recurrent evolutionary innovations in mammals.

## INTRODUCTION

*Schlafens* are a class of interferon stimulated genes that participate in immune response to viral infections (Kim and Weitzman 2022). In human, SCHLAFEN (SLFN) proteins have been shown to restrict a wide variety of viruses including HIV (Li et al. 2012; Stabell et al. 2016; Ding et al. 2022); HSV (Kim et al. 2021) and Influenza (Seong et al. 2017). While there is much diversity in their mode of action, most SLFNs also play important core cellular functions most notably by regulating cellular rRNAs and tRNAs metabolism (Pisareva et al. 2015; Murai et al. 2018; Yang et al. 2018; Chen and Kuhn 2019; Garvie et al. 2021; Metzner et al. 2022a; Metzner et al. 2022b).

In rodents and primates, these functions have been tied to unique structural features. All SLFN proteins are structurally related to eukaryotic and viral RNase E and NTPase/Helicase enzymes (Chen and Kuhn 2019; Jo and Pommier 2022). Their N-terminal regions universally encode a “SCHLAFEN core domain” combining a “SCHLAFEN box” and RNase E-related AlbA domain (Pisareva et al. 2015; Yang et al. 2018; Chen and Kuhn 2019; Garvie et al. 2021; Jo and Pommier 2022; Metzner et al. 2022a; Metzner et al. 2022b). This domain promotes SCHLAFEN dimerization creating an RNA binding pocket with endonuclease activity when catalytic residues are present (Garvie et al. 2021; Metzner et al. 2022b). This function is essential to coordinate rRNA and tRNA binding and/or cleavage upon various cellular stresses (Schwarz et al. 1998; Geserick et al. 2004; Zoppoli et al. 2012; Arslan et al. 2017; Fischietti et al. 2018; Yang et al. 2018; Yue et al. 2021; Metzner et al. 2022a).

The SCHLAFEN core domain is followed by a linker domain containing a “SWADL” motif– and with structural similarities to GTPases (Chen and Kuhn 2019). The C-terminal portion is an NTPase-related Helicase domain, sometimes referred to as P-Loop (Chen and Kuhn 2019; Jo and Pommier 2022). These domains also contribute to dimerization and interaction with other proteins (Chen and Kuhn 2019; Garvie et al. 2021; Kim et al. 2021; Jo and Pommier 2022; Metzner et al. 2022b; Nightingale et al. 2022). For some SLFN proteins, these domains coordinate nucleotide binding – *e.g.,* SLFN12 with GDP and cyclic NTPs (Chen and Kuhn 2019; Garvie et al. 2021); SLFN5 with ATP (Metzner et al. 2022a) – albeit with no hydrolysis activity on their own (Garvie et al. 2021). Moreover, the SWADL and Helicase domains have been recently shown to bind ssDNA *in vitro* (Metzner et al. 2022b), and are necessary for viral restriction *in vivo* (Li et al. 2012; Pisareva et al. 2015).

From an evolutionary stand-point, *Schlafen* genes are found across all vertebrates and likely derive from a single parental RNase E-like ortholog (Chen and Kuhn 2019). All extant vertebrate *Schlafen* genes are found within a single genomic cluster (flanked by *Unc45b* and *Pex1*2). They group into four phylogenetically distinct clades: Clade 1 relates to human *SLFN12*; Clade 2 to *SLFN5*; Clade 3 to *SLFN14;* and Clade 4 to *SLFN11* (Bustos et al. 2009). However, the total number of *Schlafen* genes found in this cluster varies greatly between species, as a result of repeated lineage-specific gain and loss events (Bustos et al. 2009). For instance, the *Schlafen* cluster encodes for one coding and two pseudogenized *Schlafens* in dogs (*Canis familiaris*); six in humans (*Homo sapiens*); and nine coding and one pseudogene in mice (*Mus musculus*) (Bustos et al. 2009; Lilue et al. 2018).

These expansions and contractions were accompanied by structural changes between and within clades (Bustos et al. 2009; Chen and Kuhn 2019; Jo and Pommier 2022). The most drastic example is the complete loss of the C-terminal helicase domain in the parental gene giving rise to Clade 1 *SLFNs* roughly 100 million years ago (Ma) followed by further shortening of the C-terminal portion during rodent evolution (Bustos et al. 2009). Finally, vertebrate *Schlafen* orthologs have been suggested to display accelerated rates of codon evolution (Bustos et al. 2009; Stabell et al. 2016; Lilue et al. 2018). In the case of primate *SLFN11*, specific codons subject to positive selection were shown to have shaped its antiviral activity most notably against HIV-1 in human cells (Stabell et al. 2016). Nevertheless, for most Schlafens, the link between sequence variation and function remains unknown.

The leading hypothesis underlying the evolutionary diversification of the *Schlafen* family is their involvement in an ongoing genetic conflict with pathogens (Bustos et al. 2009; Kim and Weitzman 2022). Genetic conflicts arise when co-evolving genetic entities engage in antagonistic interactions (Gardner and Ubeda 2017; McLaughlin and Malik 2017). This promotes rapid adaptation/counter-adaptation cycles, or evolutionary arms races, driving the expansion and positive selection of protein domains engaged in antagonism (Hurst and Werren 2001; McLaughlin and Malik 2017; Daugherty and Zanders 2019; Kuzmin et al. 2022). Considering that many primates SLFN proteins have well described antiviral functions, their rapid diversification could result from an arms race with viral proteins (Kim and Weitzman 2022). Supporting this hypothesis, human SLFN5 and SLFN11 are the targets of viral antagonisms in the context of HSV-1 and HCMV restriction respectively (Kim et al. 2021; Nightingale et al. 2022).

However, Schlafens rapid evolution might also relate to functions beyond antiviral antagonisms. This might be the case in mice where genetic mapping of a strain incompatibility locus was directly linked to the *Schlafen* gene cluster (Cohen-Tannoudji et al. 2000; Bell et al. 2006). This suggests that SLFN proteins could play an important function during reproduction and that their rapid evolution contributes to reproductive isolation, and ultimately speciation (Crespi and Nosil 2013). In this case, whether this genetic conflict relates to antiviral antagonism is unknown. More importantly, which *Schlafen* genes and which domains drive this incompatibility remains to be determined.

Here, we use a detailed phylogenomic approach to determine the evolutionary trajectories and selective forces that shaped mammalian *Schlafen* orthologs and paralogs. We trace the origins of a novel *Schlafen-like* orphan gene in vertebrates and the lineage-specific amplifications patterns of clustered *Schlafen* genes in mammalian genomes. We use in-depth codon-level selection analyses to map recent diversification events in rodent and primate genomes. We show that many recently duplicated *Schlafen* genes evolved under distinct selective forces and identify recurrent positive selection over *SLFN11*, *SLFN5* and *SLFN12*-related genes. Finally, we use crystal structures with machine learning predictions, to identify a novel class of residues subjected to positive selection mapping at the dimer interface between SLFN proteins. We suggest that this diversification shaped both antiviral and developmental functions of *Schlafen* paralogs.

## RESULTS

### Evolutionary trajectory of the *Schlafen* gene cluster and *SLFNL1* in mammals

To comprehensively study the origins and selective pressures acting on mammalian *Schlafen* orthologs, we first set out to update the evolutionary history of the *Schlafen* gene cluster using up-to-date vertebrate genomes (Bustos et al. 2009). We used TBLASTN searches with annotated SLFN proteins from Human (*Homo sapiens*), Horse (*Equus cabalus*) and Elephant (*Loxodonta africana*) to identify candidate *SLFN* genes across vertebrate and extending to several outgroups, such as Ciona (*Ciona intestinalis*) (Methods). This approach retrieved at least one putative *Schlafen* ortholog in most vertebrate genomes but none with significant homology in species outside of jawed vertebrates – i.e., in Elephant shark (*Callorhinchus milii*) but not in hagfish (*Eptatretus burgeri*) (Figure 1A). In jawless vertebrates, none of the synteny is preserved blurring further homolog identification based on assembled contigs (Figure 1A).

**Figure 1.**
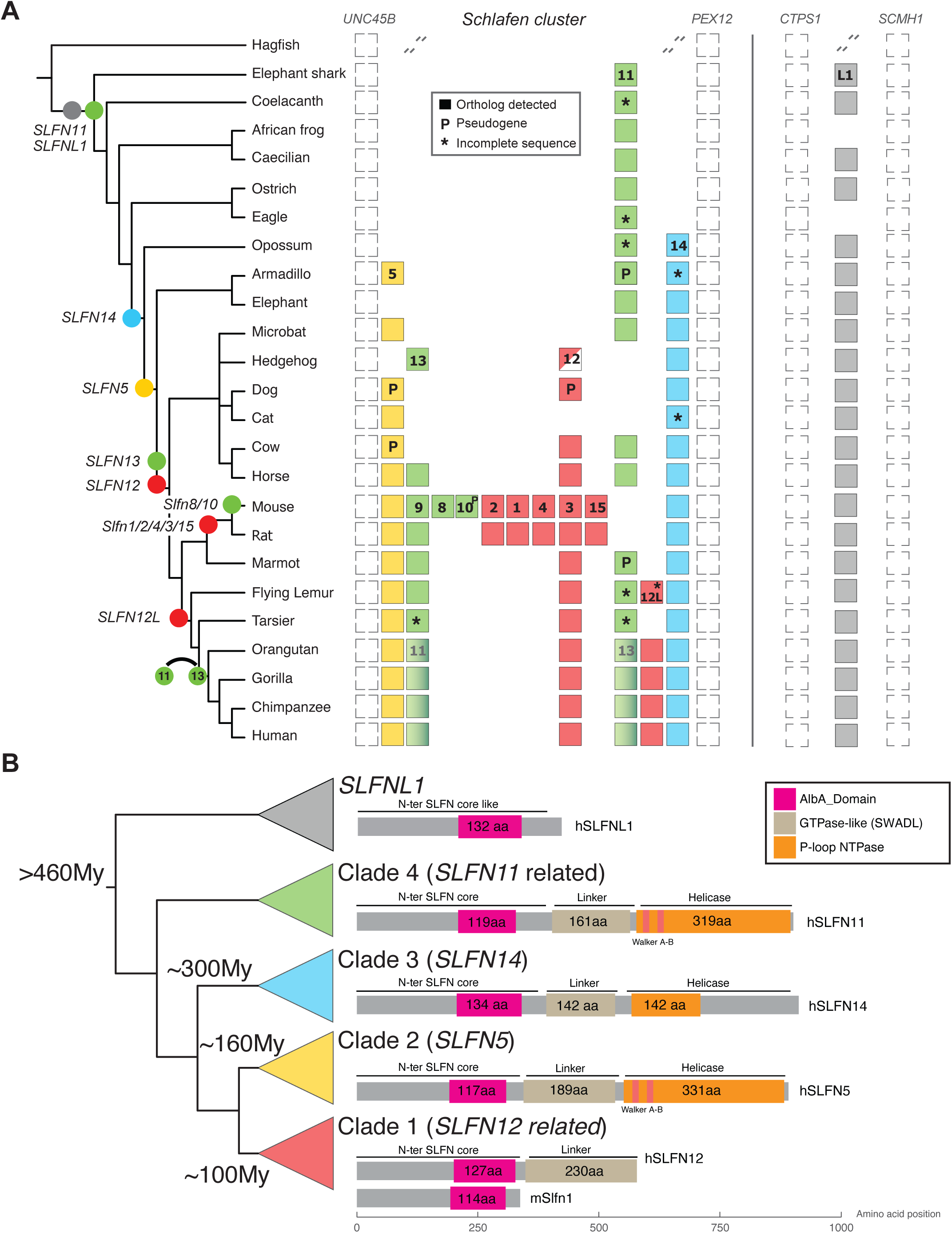
Evolutionary history of *Schlafen* genes in Vertebrates. (**A**) Schematic representation of genomic loci encoding clustered *Schlafen (SLFN)* and *Schlafen Like 1 (SLFNL1)* genes in selected species of vertebrates. Filled circles denote major gene birth events along the accepted species phylogeny. For each clade, presence of *Schlafen* and *SLFNL1* loci are shown with filled boxes with the associated *Schlafen* gene family number shown for the top instance. Empty spaces indicate that no putative ortholog was found in the assembly. Genes flanking each locus are shown with empty boxes and dashed backslashes indicate a break in synteny. “P” denotes pseudogenes and “*” incomplete sequences. A switch in *SLFN13* and *SLFN11* is shown at the base of primates. Incomplete lineage sorting of Hedgehog putative *Slfn12* is shown with a half-filled box. (**B**) Inferred evolutionary history of *Schlafen* and *SLFNL1* clades based on phylogenetic and syntenic analyses. Clades encoding for *SLFN12* (red), *SLFN5* (yellow), *SLFN14* (blue), *SLFN11* (green) and *SLFNL1* (gray) are shown with their estimated date of birth. For each clade, representative secondary structures of human or mouse protein are shown with their associated domains mapped relative to the starting amino-acid (bottom X-axis).

To identify specific orthologs, we first analyzed coding sequences (CDS) syntenic location on existing genome assemblies or using manual contig annotation (see Methods). We found that most hits mapped back to the expected *Schlafen* gene cluster flanked by the *Unc45b* and *Pex12* genes (Bustos et al. 2009) (Figure 1A). However, one hit consistently mapped outside of the known *Schlafen* cluster between the *Ctps1* and *Scmh1* genes (Figure 1A). This locus codes for a poorly characterized *Schlafen-*related gene found in most vertebrate genomes, named *SLFNL1* in human (RefSeq accession NM_144990). While SLFNL1 predicted protein shares less than 10% identity with other SLFNs, it nonetheless contains a SCHLAFEN AlbA domain with conserved catalytic residues indicative of potential RNAse E functionality (Supplemental Fig. S1). Thus, our syntenic analysis suggests that the last common ancestor to all jawed vertebrates likely encoded two potentially enzymatically active *Schlafen* genes, one within the cluster and one orphan related to *SLFNL1*.

Having identified the genomic location of all *Schlafen* homologs, we wished to uncover their evolutionary relationships to finalize ortholog assignment based on shared ancestry. Considering the large time span of our analysis – ∼500My (Million years) – we used predicted SLFN proteins to run maximum likelihood (ML) phylogenetic analyses (see Methods). This identified five distinct *Schlafen* monophyletic clades: the four previously reported *Schlafen* gene clades with the addition of one novel clade related to *SLFNL1* (Figure 1B, Supplemental Fig. S2 & Supplemental Table S1). Increased divergence did not allow proper grouping of non-mammalian vertebrate SLFN11 with their orthologs (Supplemental Fig. S2). Yet, these homologs code for all the key residues and domains characteristic of this family, supporting their shared ancestry (Supplemental Data S1). Finally, within mammals, the hedgehog putative *SLFN12* ortholog grouped outside of both *SLFN5* and *SLFN12* clades (Supplemental Fig. S2). Indeed, while its N-terminal portion is characteristic of other *SLFN12*, it appears to have acquired an extended C-terminal distantly related to *SLFN5* (Supplemental Fig. S3).

Combining our genomic and phylogenetic analyses, we made several interesting observations. First, the last common ancestor to all jawed-vertebrate likely encoded for one orthologs of *SLFN11* (clade 4) located in the *Schlafen* gene cluster, and one orphan *SLFNL1* (Figure 1A). These two *Schlafens* remained in most vertebrate genomes, apart from Carnivores that lost *SLFN11*-related genes, and some amphibians and birds that independently lost *SLFNL1* (Figure 1A). Second, the *Schlafen* gene cluster expanded in mammals with the successive birth of *SLFN14* (∼300Ma), *SLFN5* (∼160Ma) and more recently *Slfn12*-related genes (∼100Ma) (Figure 1).

Following these birth events, *SLFN14* remained intact in all mammalian genomes investigated, while all other clustered *Schlafens* underwent lineage-specific loss/amplification events (Figure 1A). A few examples include: (1) the duplication of a parental *SLFN11* into *SLFN13/11* paralogs following the split of Atlantogenates (*i.e.,* elephants & armadillos) from the rest of placental mammals; (2) the amplification burst of *Slfn12*-related paralogs following the split between marmots and other rodents (discussed in detail below); or (3) the birth of *SLFN12L* in the last common ancestor of primates and dermoptera (Bustos et al. 2009). In sum, our phylogenomics approach identified a novel orphan *Schlafen*-related gene that likely arose in vertebrates, and confidently assigned orthology between *Schlafen* cluster paralogs allowing for accurate natural selection studies.

### Diversifying selection shapes *Schlafen* genes in Primates

As seen for other immune-related genes, SLFN antiviral functions are expected to drive recurrent evolutionary diversification, or positive selection, in response to pathogens (McLaughlin and Malik 2017). While this has been demonstrated for primates’ SLFN11 N-terminal region (Stabell et al. 2016), whether similar selective forces also shape other paralogs remains unexplored. To address these shortcomings, we retrieved and annotated *Schlafen* orthologs in simian primates – which includes new world monkeys, old world monkeys and great apes – and used Tarsiers as an outgroup. This group is well suited for natural selection analysis as it spans ∼60My without saturation of synonymous mutation rates over 20 annotated genomes (Perelman et al. 2011).

We found that all primates have an intact *Schlafen* cluster encoding orthologs of *SLFN5, SLFN11, SLFN12, SLFN13, SLFN12L, SLFN14,* in addition to an intact *SLFNL1* (Figure 2, Supplemental Fig. S2 and Supplemental Table S1). However, following the split with tarsiers, an inversion switched *SLFN11* and *SLFN13* locations but left the rest of the cluster unaffected (Figure 1A, and Supplemental Table S1). Closer investigation of the locus could not resolve the genetic nature of this event since *SLFN11* and *SLFN13* surrounding regions as well as the other *Schlafen* loci in the cluster kept their parental orientation. However, primate *Schlafen* orthologies were well supported by maximum likelihood phylogenies (Figure 2 and Supplemental Table S1).

**Figure 2.**
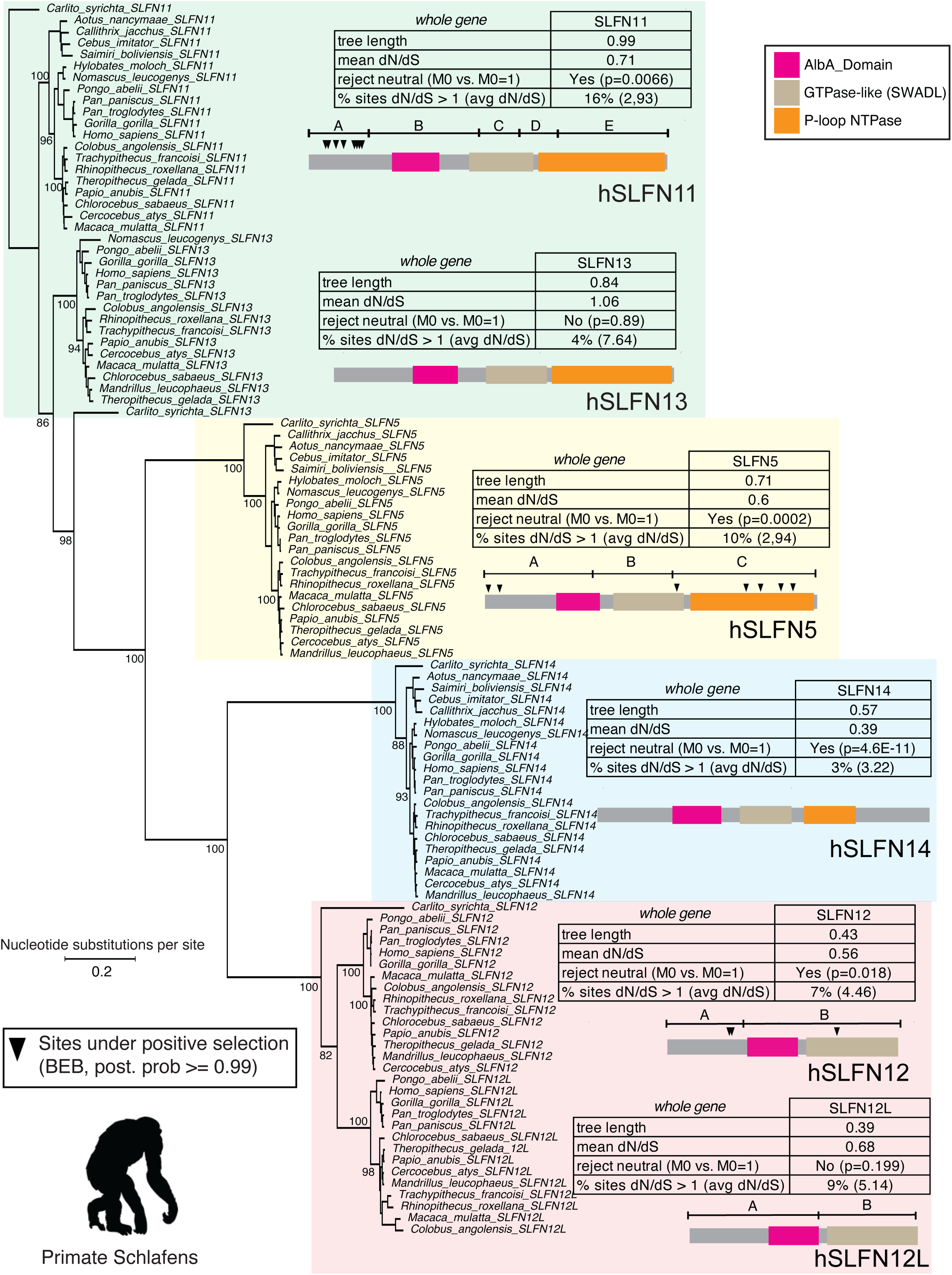
Rapid evolution of *Schlafen* genes in primate genomes. Maximum likelihood phylogeny of all identified Schlafen coding sequences in simian primates. The tree is artificially rooted on Tarsier *SLFN11*. Only major bootstrap support values >50% are shown. For each highlighted clade, a summary table of selection analyses on whole CDS are shown (“whole gene”). Sites evolving under positive selection (PAML, BEB posterior probability >99%) are mapped with arrowheads under each putative recombination segments detected by GARD. Chimpanzee silhouette is from https://www.phylopic.org.

To determine the strength and direction of natural selection, we measured synonymous (dS) and non-synonymous (dN) codons substitution rates across codon-aligned predicted CDS (see Methods). We first used likelihood approaches to test if modeling dN/dS across the whole gene showed significant deviation from neutral evolution (dN/dS close to 1). While most *Schlafen* genes showed strong support for natural selection (Figure 2 and Supplemental Table S2), we failed to reject neutral evolution for *SLFN13* (mean dN/dS=1.06, see Methods) and *SLFN12L* (dN/dS=0.68). Weak signatures of selection could indicate that these orthologs accumulate mutations without measurable fitness consequences and might be on their way to becoming non-functional in primates.

Of those *Schlafens* under selection, *SLFN14* had the lowest overall dN/dS (0.39) likely reflecting predominant purifying selection over core housekeeping functions (Figure 2 and Supplemental Table S2) (Pisareva et al. 2015; Seong et al. 2017). For other paralogs, we turned to codon-level dN/dS likelihood-based modeling to determine which residues and domains might be subject to positive selection (see Methods). Considering the clustering of *Schlafen* genes, they could experience increased recombination rates sometimes leading to pervasive gene conversion events within or between paralogs (Mitchell et al. 2015; Daugherty et al. 2016; Daugherty and Zanders 2019; Molaro et al. 2020). These events can alter local substitution rates over specific CDS portions and thus blur codon-level selection analyses that model homogeneous dN/dS across whole CDS. To account for this potential caveat, we used GARD to predict putative recombination segments (see Methods) (Kosakovsky Pond et al. 2006). When detected, we performed separate selection analyses on each predicted segment, otherwise we used whole CDS (Figure 2, Supplemental Table S2).

This approach identified eight codons under recurrent positive selection clustered over the N-terminal end of SLNF11 (segment A, BEB>0.99, Figure 2, Supplemental Table S2). Of these positively selected codons, seven were directly overlapping with a prior study (M47, Q79, C102, Y133, E137, V140, S142) attesting of the robustness of our analysis (Stabell et al. 2016). Using the recently published SLFN11 crystal structure (Metzner et al. 2022b), we found that five of these sites mapped inside the RNase E binding pocket and are thus likely to affect SLFN11 enzymatic function (Supplemental Fig. S4).

The remaining three residues mapped toward the surface of the protein. However, unlike what was reported before, we found that two of these residues (Q79 and E137) located at the direct contact interface between SLFN11 molecules in the crystalized homodimer (Figure 3A). Finding 2 out of 8 positively selected residues (1/4) at this interface is highly unlikely since there are only 37/901 (∼1/24) total residues within <4Å between the two monomers (chi-square test p-value = 0,011) (Figure 3A). Thus, this indicates the SCHLAFEN contact interface is enriched for diversifying residues. Interestingly, in the case of codon Q79, its side chain in one monomer contacts a region neighboring two other rapidly evolving codon in the second monomer (V140 and S142, Supplemental Fig. 4). This last observation further suggests that intra-SLFN molecular interactions might be subject to ongoing diversification in primates.

**Figure 3.**
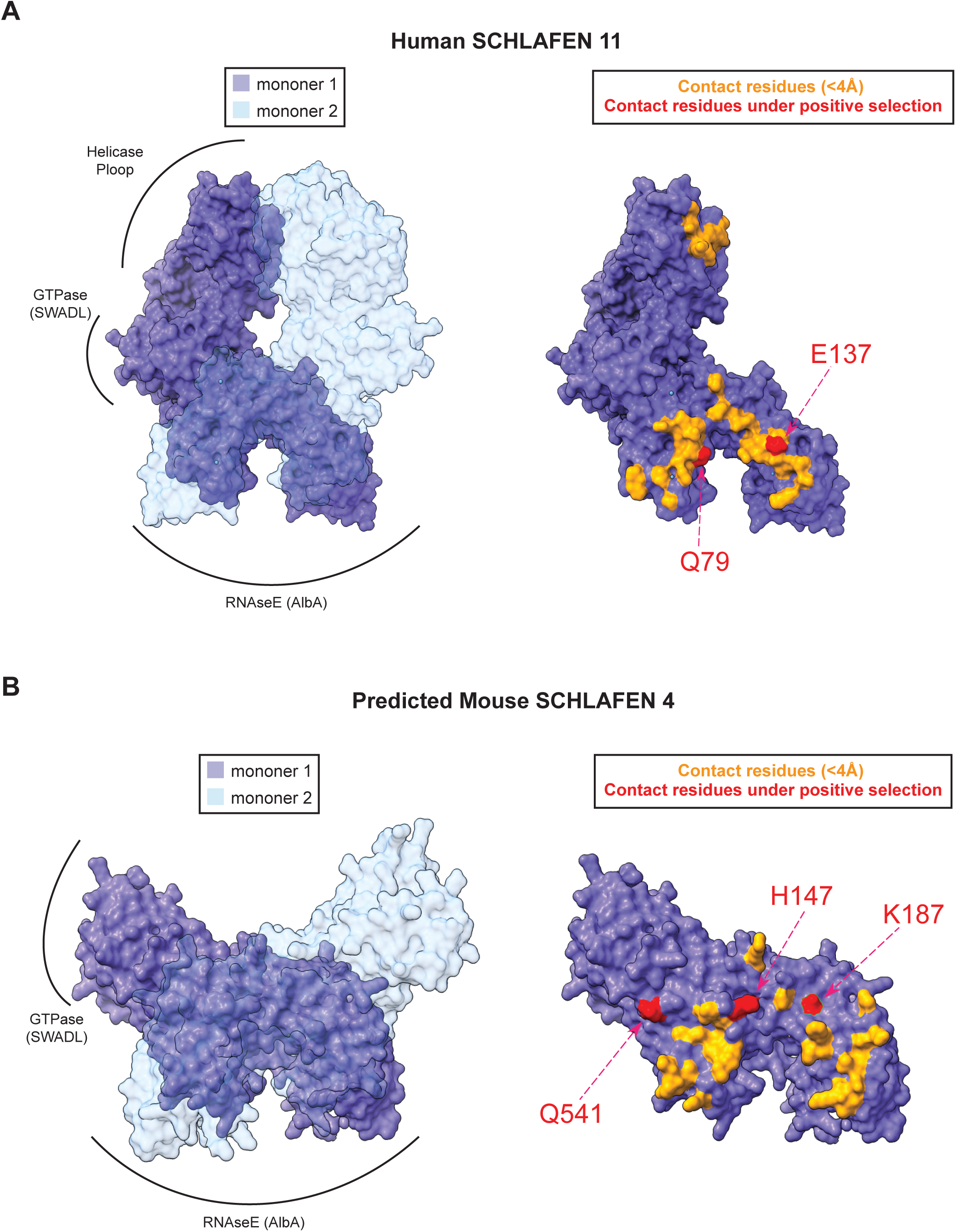
Positive selection over SCHLAFEN homodimer contact residues. (**A**) Surface representation of crystalized Human SLFN11 homodimer (PDB: 7ZEP). Major domain localizations are shown along monomer 1 (purple structure). Monomer 2 is shown in transparency (light blue). Residues within 4Å between monomer 1 and 2 (contact residues) are highlighted in yellow onto monomer 1 (right panel). Contact residues under positive selection are shown in red with their relative position in the human protein. (**B**) Same as (A) but for Alphafold2 predicted mouse SLFN4 homodimer. Positions are relative to the mouse protein.

Beside SLFN11, our analysis also uncovered several interesting new cases of positive selection in primates. First, we identified seven positively selected sites in *SLFN5* (Figure 2, Supplemental Table S2). Suggestive of similar selective pressure than those acting on *SLFN11,* two of these rapidly evolving codons mapped to its N-terminal portion (segment A, Figure 2, Supplemental Fig. S5). Since SLFN5 lost its RNase activity, the functional significance of these changes is unknown (Metzner et al. 2022a). Outside of the N-terminus, we found four diversifying residues within the C-terminal Helicase domain (P-loop NTPase, segment C, Figure 2, Supplemental Table S2). SLFN5 helicase is implicated in both viral and retrotransposon activity and is the direct target of the viral antagonist ICP0 encoded by HSV-1 (Kim et al. 2021; Ding et al. 2023). Of note, diversification over the helicase domain was not found in other *Schlafens*.

Second, we identified three positively selected residues over the 5’ segment of *SLFN12* (Figure 2, Supplemental Table S2). This contrasts with its duplicate gene *SLFN12L* that displays low overall level of selection (see beginning of section). Mapping these residues over the SLFN12 structure did not show specific enrichment in the RNase domain or at the interface between monomers, raising questions on the selective forces driving SLFN12 diversification (Supplemental Fig. S6). Nevertheless, this finding indicates that *SLFN12* and *12L* have been subject to different selective forces following their duplication ∼85Ma and likely carry-out independent functions in today genomes.

In sum, our approach identified recurrent diversifying selection over the primate *Schlafen* genes *SLFN11, 5* and *12*. While many rapidly evolving residues map to known domains with antiviral functions, we report novel sites that locate at their dimerizing interfaces. This observation suggests that intra– and/or inter-SLFN interactions have been under strong selection during primate evolution.

### Birth of *Slfn12-* and *Slfn13*-related genes in rodents

Compared to primates, mouse *Schlafen* genes received less attention. This partially stems from the complex structure of *Schlafen* paralogs within the mouse cluster (Lilue et al. 2018). Indeed, in addition to encoding *Slfn14* and *Slfn5,* and unlike any other mammals, the mouse cluster codes for multiple *Slfn12* and *Slfn13*-related genes with uncharacterized phylogenies (Figure 1A) (Schwarz et al. 1998; Bell et al. 2006; Bustos et al. 2009; Lilue et al. 2018). We used our in-depth phylogenomics approach to map these evolutionary transitions over 22 species spanning ∼70My of rodent speciation (Kumar et al. 2017; Swanson et al. 2019).

We found that the amplification of *Slfn12*-related genes occurred in three successive duplication rounds (Figure 4A, Supplemental Table S1). First, an event giving birth to one GTPase-less *Slfn12* and two full-length paralogs (*Slfn3* and *Slfn4*) occurred following the split between degus and other rodents ∼50Ma (Figure 4A). A second round of duplication led to the formation of two additional GTPase-less paralogs (*Slfn1* and *Slfn2)* in the last common ancestor of muroids rodents (∼43Ma, Figure 4A). One final duplication occurred in the last common ancestor of mice and rats (∼16Ma) and gave birth to a third full length gene *Slfn15* (Figure 4A). We note that this gene was previously annotated as *GM11427*. Although maximum likelihood phylogenies only partially resolved each paralog family, likely due to pervasive gene conversion (see next section), orthology was well supported using syntenic gene structure within contigs or chromosomes (Supplemental Fig. S7).

**Figure 4.**
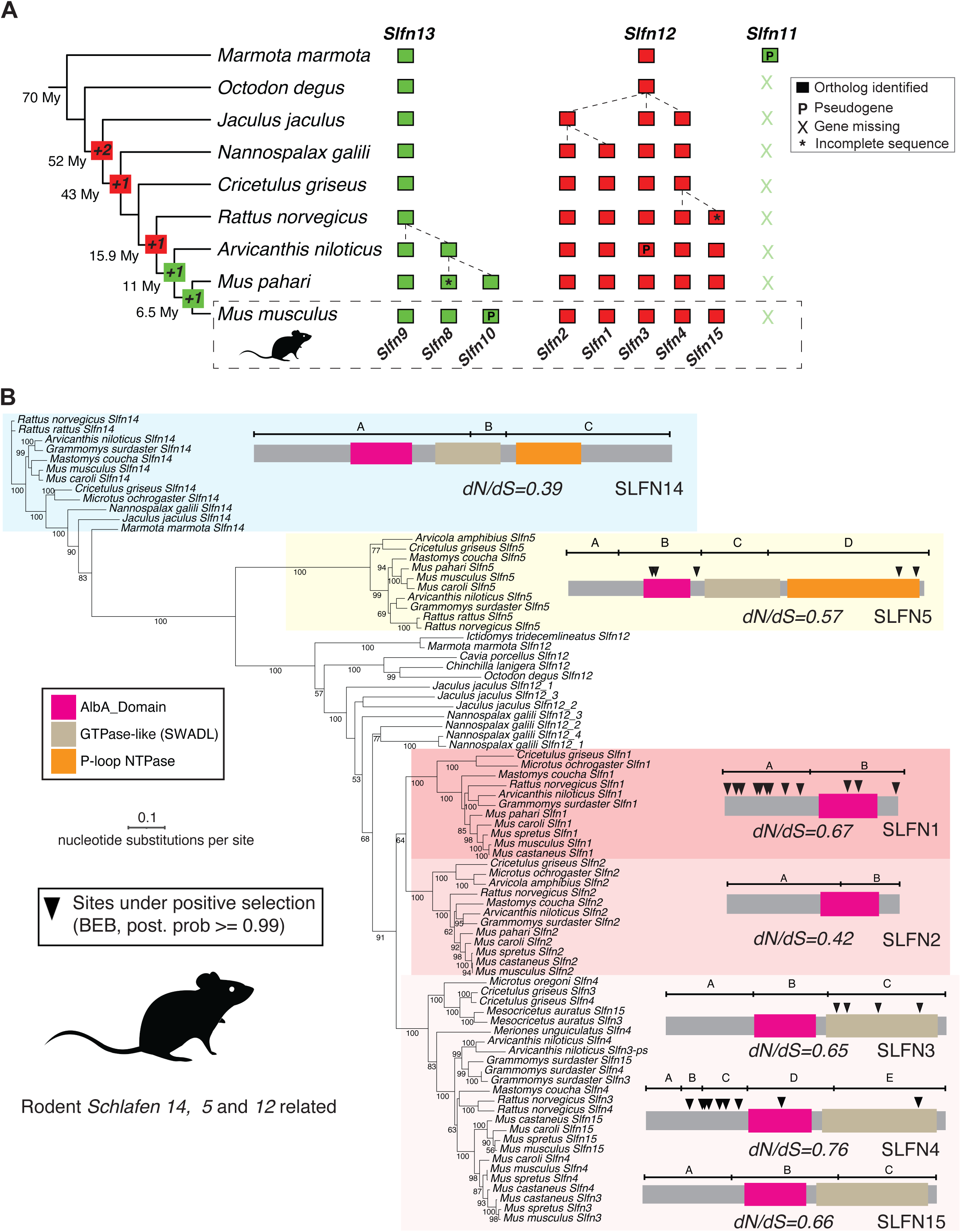
Amplification and rapid evolution of *Schlafen* genes in rodent genomes. (**A**) Summary of *Slfn12* and *Slfn13* duplication events along the rodent species tree with estimated divergence times. Identified orthologs are shown with filled boxes. Gene names are shown relative to the Marmot (top) and mouse (bottom) genomes. Pseudogenes are indicated with “P” and incomplete sequences with “*”. *Slfn11* is missing from most genomes following the split with marmots (denoted with “X”). (**B**) Maximum likelihood phylogeny of rodent *Slfn14, Slfn5* and *Slfn12*-related coding sequences. Bootstrap support values >50% are indicated. Average dN/dS for each *Slfn* clade are indicated below a schematic protein secondary structure. Codon evolving under positive selection (PAML, BEB posterior probability >99%) are mapped with arrowheads under each putative recombination segments detected by GARD. Mouse silhouette is from https://www.phylopic.org.

*Slfn13* related genes showed a much more recent evolutionary origin, with amplification beginning after the split with rats (∼16Ma) and continuing in *Mus* species (Figure 4A). The first duplication gave birth to *Slfn9* and the ancestor of *Slfn8/10,* which subsequently duplicated in the last common ancestor between *Mus pahari* and *Mus musculus* (Figure 4A). This event was quickly followed by a pseudogenization of *Slfn10* in many *Mus musculus domesticus* strains (Supplemental Fig. S8). Phylogenetic analyses, and when possible synteny, supported that *Slfn13* was the parental gene subject to these duplications (Supplemental Fig. S9). This is further supported by the fact that *Slfn11* pseudogenized early during rodent evolution (Figure 4A).

In sum, we were able to trace the origins of clustered *Schlafen* genes in rodents (leading to ten genes in mouse compared to six in human). We identified a *Slfn12* duplication event accompanied by full domain loss giving rise to GTPase-less clustered *Schlafens* unique to rodents. These have been retained for at least 50My, indicative of putative functions. We also found an ancient loss of *Slfn11* and, more recently, a burst of *Slfn13* duplications. These likely reflects recent switches in selective pressures since these two genes have been stable during most of mammalian evolution. Next, we explored whether these events were accompanied by signatures of natural selection indicative of recent functional innovations.

### Concerted evolution affecting GTPase-containing *Slfn12* duplicates

To detect putative recombination in rodent *Schlafen* orthologs we used GARD analysis as described in the previous section. This detected several putative recombination segments in all rodent *Schlafen* clades (Supplemental Table S3). We hypothesized this might be due to concerted evolution as many *Schlafen* orthologs failed to segregate according to their orthology in whole CDS phylogenies (Figure 4B).

Indeed, outside of *Mus* subspecies, *Slfn3*, *4* and *15* paralogs grouped within species and not with their orthologs (Supplemental Fig. S10A). Interestingly, this was not the case for GTPase-less *Slfn1* and *Slfn2* orthologs (Supplemental Fig. S10A). This pattern was also observed in distant species with “unresolved” *Slfn12*-related paralogs (*e.g., Nannospalax galili* or *Jaculus jaculus,* Supplemental Fig. S10A). To test whether this was the result of direct gene conversion between paralogs, we compared the genomic identities of the encoding loci (Supplemental Fig. 10). This revealed extensive nucleotide identities in the introns and flanking regions of both *Slfn12_4* and *_1* in *Nannosplalax* galili (Supplemental Fig. S10B) and *Slfn3* and *4* in *Cricetulus griseus* (Supplemental Fig. S10C). Together these results indicate that GTPase-containing *Slfn12* paralogs have been subject to repeated gene conversion since their birth 50Ma (Figure 4A). The selective forces preventing these events from occurring between GTPase-less paralogs, of about the same age, remains unknown.

### Novel signatures of positive selection amongst rodent *Schlafen* genes

For our natural selection analyses we used individual recombination segments of coding CDS (see Methods and previous section). As observed for their primate orthologs, we found that *Slfn14* had the lowest overall dN/dS of all *Schlafen* clades, suggesting purifying selection (Figure 4B, Supplemental Table S3). *Slfn5* orthologs showed pervasive positive selection with several positively selected codons mapping to the helicase domain (Figure 4B, Supplemental Table S3, Supplemental Fig. S11). Thus, *Slfn5* and *Slfn14* are subject to similar selective forces in primates and rodents despite ∼87My of divergence. This suggests deeply conserved functions for these two *Schlafen* orthologs including in the context of viral restriction.

*Slfn12*-related orthologs displayed more complex selective signatures. First, we found that the two GTPase-less paralogs, *Slfn1* and *Slfn2*, evolved under different selective constraints. While *Slfn2* showed signatures of purifying selection (overall dN/dS=0.42, Supplemental Table S3), *Slfn1* orthologs displayed evidence of diversification (dN/dS=0.67, Supplemental Table S3). Codon-level selection analyses revealed that many sites accumulated over Slfn1 N-terminus with two overlapping the RNase E/Alba domain (Figure 4B, Supplemental Table S3). We currently lack a crystal structure of this protein, and its sequence divergence prevented us from mapping these sites on human SLFN12. However, we used AlphaFold2 to predict SLFN1 monomers and dimers (see Methods). While none of the predicted dimers passed our significance threshold, predicted monomers had the expected SLFN structural features (see Methods and Supplemental Fig. S12). When we mapped rapidly evolving sites over this prediction, we found that half were located on the side of the monomer engaged in dimerization in other SCHLAFEN orthologs (L66, S67, S85, P87, V109, N143, Supplemental Fig. S12). This echoes our findings for primate SLFN proteins.

Second, of all rodent GTPase encoding *Slfn12* paralogs, *Slfn3* showed the strongest signature of selection including positive selection over codons belonging to the GTPase domain (Figure 4B, Supplemental Table S3, Supplemental Fig. S13). This contrasts with *Slfn4* orthologs which, albeit weaker evidence for overall selection, accumulated rapidly evolving codons at their N-terminus (Supplemental Table S3, Supplemental Fig. S13). AlphaFold2 predictions of SLFN3 and SLFN4 homodimers matched known SLFN structural features (Supplemental Fig. S13 and Supplemental Fig. S14) (Garvie et al. 2021; Metzner et al. 2022b). Again here, rapidly evolving SLFN4 N-terminal associated sites were enriched at the dimer interface (Figure 3B). Indeed, 3 out of 9 positively selected residues (1/3) locate at this interface, compared to 29 out of 602 (∼1/21) total residues within <4Å between the two monomers (chi-square test p-value = 0,0013) (Figure 3A) Finally, we failed to detect evidence for positive selection over *Slfn15* orthologs suggesting they might be evolving primarily under purifying selection, albeit with an average dN/dS=0.6 (Supplemental Table S3). The distinct domain distribution of sites under positive selection between *Slfn3, Slfn4* and *Slfn1*, suggests that their parental *Slfn12* gene might have carried multiple functions subject to rapid diversification.

Considering that most *Slfn13* duplicates arose in *Mus* sub-species, we extended our selection analyses to *Schlafen* orthologs across *Mus musculus musculus* and *domesticus* strains (see Methods). We used whole coding sequences to run McDonald-Kreitman (MK) testing to compare the rates of synonymous to non-synonymous substitutions within *Mus musculus* strains (polymorphisms) to those fixed between species using *Mus Spretus* or *Mus Caroli* as outgroups. This identified *Slfn8* and *Slfn9*, but not *Slfn10*, as evolving under positive selection (Supplemental Table S4). Furthermore, we found whole gene diversifying selection amongst rodent *Slfn13* orthologs (Supplemental Table S4). This suggests that recent *Slfn13* duplicates might share similar selective pressures with their parental gene. Unfortunately, PAML analyses lacked the phylogenetic depth to significantly identify the residues most likely subject to this diversification on rodent *Slfn13* segments or full-length sequences in mouse strains (Supplemental Table S4). Nevertheless, this result is in line with prior reports in mice (Lilue et al. 2018), and indicates that these recent duplicates have continued to diversify their parental gene function.

In sum, we detected novel events of diversifying selection over recent rodent-specific *Slfn12* duplicates indicating that the birth of *Slfn1/2* and *Slfn3/4* paralogs was accompanied by repeated sub-functionalization. As reported in primates, many rapidly evolving sites accumulated at predicted SLFN interaction surfaces suggesting that SCHLAFEN dimerization is under strong selection in rodents and further supporting its functional importance across mammals.

## DISCUSSION

Genetic conflicts are key drivers of genome evolution. Most notably, they accelerate genetic innovations over short evolutionary timescales in the form of positive selection and gene family expansion (McLaughlin and Malik 2017). Here, we detected unprecedented signatures of evolutionary diversification of clustered *Schlafen* genes in primate and rodent genomes. *Schlafen* genes engage in at least two genetic conflicts: host/virus restriction in primates; and possibly hybrid incompatibilities in rodents (Cohen-Tannoudji et al. 2000; Bell et al. 2006; Bustos et al. 2009; Lilue et al. 2018; Kim and Weitzman 2022). In the following section, we discuss our findings in the context of these functions.

Building from pioneering genomics studies, our depth phylogenomic analyses dated the birth of clustered *Schlafens* to the last common ancestor to all jawed vertebrates, roughly 450Ma (Hedges et al. 2015). We found that a *Slfn11*-like gene was the parental gene of the cluster. SLFN11 is able to restrict retroviruses in several mammalian species suggesting deep antiviral functions (Lin et al. 2016; Stabell et al. 2016; Kim and Weitzman 2022). It is thus tempting to speculate that selective pressures from emerging viruses contributed to *Schlafen* cluster diversification early during vertebrate radiation. This diversification intensified in mammals with the successive birth of *Slfn14*, *Slfn5*, *Slfn13* and finally *Slfn12*. Some of these families, *e.g., Slfn13* and *Slfn12* (Basson et al. 2018; Yang et al. 2018; Garvie et al. 2021; Yue et al. 2021), function beyond pathogen restriction indicative of continued sub-functionalization. Our finding of diverging, and sometimes opposite, selective forces over recent amplifications – *SLFN12/12L* and *SLFN11/13* in primates, *Slfn12* duplicates in rodents – further supports this conclusion.

Our analysis uncovered that one non-clustered *Schlafen* gene, *SLFNL1*, is also ancestral in vertebrate genomes. SLFNL1 has yet to be functionally characterized and shares only that RNase E domain with other SLFNs such as SLFN11. Recently, structurally similar SLFN-like proteins have been shown to help piRNA processing in C.elegans (Podvalnaya et al. 2023). Thus, it is possible that orphan genes such as *SLFNL1* might carry out ancestral functions in RNA biogenesis, perhaps related to retrotransposon control (Bartonicek et al. 2022; Ding et al. 2023; Podvalnaya et al. 2023). If so, clustered SLFN antiviral functions might have arose following the addition of regulatory domains such as GTPase and Helicase domains to the “SCHLAFEN core”.

One of the major findings of our study is the pervasive signature of positive selection over both rodent and primate *SLFNs*. Supporting an ongoing genetic conflict with viruses, we found that rapidly evolving codons in primate *SLFN11* and *SLFN5* mapped to known protein domains engaged in viral restriction or subjected to viral antagonism (Stabell et al. 2016; Kim et al. 2021). For SLFN11, these fall within its RNase pocket. While it is currently unknown whether the SLFN11 viral antagonist, HCMV protein RL1, specifically targets this domain. Yet, this domain is essential for SLFN11 function during HIV and HCMV restriction (Stabell et al. 2016; Nightingale et al. 2022). Since we found similar signatures in rodent *Slfn1* and *Slfn4*, it would be interesting to investigate whether RNase activity is also engaged in viral restriction in these species.

SLFN5 positive selection mapped to the Helicase domain targeted by HSV-1 protein ICP0 supporting a putative antagonistic co-evolution between SLFN5 and HSV-1 (Kim et al. 2021). Again here, a similar signature is found in rodent *Slfn5* highlighting shared evolutionary constraints. However, this domain is also shown to bind single stranded DNA *in vitro* and is required for retrotransposon restriction in human cells (Metzner et al. 2022b; Ding et al. 2023). Thus, we cannot exclude that additional molecular antagonisms might also drive SLFN5 rapid evolution. Nevertheless, the diverse immune functions of *Schlafens* familiy members beyond those reported for Slfn11 and Slfn5 unanimously points to host/pathogens arms races being important drivers of their rapid evolution. Thus, future functional investigations of the positively selected sites reported here – most notably over *Slfn12*-related genes – should uncover novel essential interactions shaping mammalian immune responses.

In addition to expected host/pathogen interactions, we uncovered a novel class of rapidly evolving residues located at the interface between two SLFN proteins. So far, we are unaware of studies that directly investigated the consequences of altering SLFN protein contact interfaces *in vivo* (Jo and Pommier 2022; Kim and Weitzman 2022). Yet, biochemical work on SLFN11 and SLFN12 demonstrated the importance of homodimerization for RNA binding and cleavage likely underlying core SCHLAFEN functions (Garvie et al. 2021; Metzner et al. 2022b). Thus, our results raise several interesting hypotheses regarding the functional consequences of contact site rapid evolution.

First, we posit that contact site diversification shapes inter– or intra-SLFN compatibilities in the context of rapid paralog expansion (Kuzmin et al. 2022). In this case, the dimerizing surface might be under strong selective pressure to avoid “toxic” inter-paralog dimerization, while keeping intra-ortholog interactions untouched. This type of intragenomic conflict might be particularly important in the context of SLFN functions in hybrid incompatibilities in mice (Cohen-Tannoudji et al. 2000; Bell et al. 2006). This is supported by the recurrent shifts in selection regime we report for rodent *Slfn12* or *Slfn13* duplicates, and by the fact that rapidly evolving residues “face” each other in crystalized or predicted homodimers (SLFN11, SLFN4 and SLFN1). Further supporting this model is the observation that SLFN surfaces known to engage with other proteins do not display signatures of diversification (Arslan et al. 2017; Basson et al. 2018; Garvie et al. 2021).

Another, non-mutually exclusive hypothesis would be that other proteins interfere with dimerization via binding to this surface. One likely candidate is the orthopoxvirus protein p26 (v-Slfn poxin) that appearS to have acquired a SCHLAFEN-like domain from rodents (Gubser et al. 2007; Bustos et al. 2009). While recent work demonstrated that p26 is key to poxvirus virulence, whether it directly interacts with host Schlafens remains unknown (Eaglesham et al. 2020; Hernaez et al. 2020). In both scenario, adaption cycles against “toxic” interactions are expected to trigger evolutionary arms races driving the rapid diversification of *Schlafen* genes.

In conclusion, recurrent evolutionary innovations of the *Schlafen* gene cluster illustrate genetics conflicts’ profound consequences on genome evolution. Schlafen paralogs have been faced with the challenge of balancing rapid antiviral adaptation with their core cellular functions. Understanding how these evolutionary negotiations influenced their molecular functions should guide future investigations in the context of development, immunity and speciation.

## METHODS

### Identification of vertebrate Schlafen homologs

We performed TBLASTN searches (Altschul et al. 1990) on the NCBI nonredundant nucleotide database of 25 vertebrate genomes (see queried genomes) using translated coding sequences as queries from: (**1**) human *SLFN5* (NM_144975.4), *SLFN11* (NM_152270.4), *SLFN12* (NM_001289009.2), *SLFN14* (NM_001129820.2), *SLFNL1* (XM_011540945.2); (**2**) horse *SLFN5* (XM_005597565), *SLFN12* (XM_023653702.1), *SLFN11* (XM_005597562.3), *SLFN14* (XM_023653004.1); and (**3**) Elephant *SLFN11 (*XM_010594583.2), *SLFN14* (XM_023553377.1). We also used Elephant shark and Coelacanth *SLFN11* (XM_007906163.1; XM_014497680.1) or *SLFNL1* (XM_0078948981.1; XM_014486747.1) as queries to investigate jawless vertebrates and bilaterians. For each search, we collected the longest isoform of hits with more than 40% identity. To search for putative specie-specific homologs, we re-ran TBLASTN searches using within species homologs as queries. Finally, we performed specific TBLASTN searches on whole genome shotgun databases. A gene was considered missing when no homologous sequence (coding or non-coding) could be identified using any of these three approaches. All identified *Schlafen* sequences are available in Supplemental Table S1, raw alignment of all identified vertebrate SCHLAFENS protein in Supplemental Data S1.

### Pseudogenes and incomplete sequences

Schlafen orthologs were considered pseudogenes if we detected stop codons substitutions in their open reading frame or frame shifting mutations disrupting all recognizable homologies in predicted proteins. To detect these events, we used codon alignments with: mouse as reference for rodents; human for primates; and any closely related ortholog for other vertebrates. Partial sequences with missing exons or domains were classified as incomplete.

### Identification of Primates Schlafens

To identify *SCHLAFEN* homologs in primates, we performed TBLASTN on the NCBI nonredundant nucleotide database in primate genomes (taxid: 9443) using all the translated sequences of human *Schlafen* genes: human *SLFN5* (NM_144975.4), *SLFN11* (NM_152270.4), SLFN13 (NM_144682.6), *SLFN12* (NM_001289009.2), SLFN12L (XM_024450521.1), *SLFN14* (NM_001129820.2), and *SLFNL1* (XM_011540945.2). We collected longest isoforms of hits with more than 60 % identity. Raw alignment of all identified primate *SCHLAFEN* coding sequences is available in Supplemental Data S2.

### Identification of Rodents Schlafens

We used mouse translated sequences to search homologs in rodent species (taxid: 9989) with queries from: *Slfn1* (NM_0114072), *Slfn2* (NM_011408.1), *Slfn3* (NM_001302559.1), *Slfn4* (NM_001302559.1), *Slfn5* (XM_017314615.1), *Slfn8* (NM_1815454.4), *Slfn9* (XM_006533235), *Slfn10-ps* (NR_073523.1), *Slfn14* (NM_001166028.1) and *Slfn15* (refined assembly of XM_487182). For mouse strains, we used TBLASN searches in whole genome assemblies from 13 strains (Ferraj et al. 2023) to identify individual exons and assemble them according to their relative position and orientation in contigs. Raw alignment of all identified rodent *Schlafen* coding sequences is available in Supplemental Data S3 and mouse strains *Slfn13*-related Supplemental Data S4.

### Queried Genomes

Vertebrates: *Homo sapiens* (UCSC hg38), *Pan troglodytes* (UCSC panTro6), *Gorilla gorilla* (UCSC gorGor6), *Pongo abelii* (UCSC ponAbe3), *Carlito syrichta* (UCSC tarSyr2), *Galeopterus variegatus* (UCSC galVar1), *Marmota marmota* (NCBI GCF_001458135.2), *Rattus norvegicus* (UCSC Rn6), *Mus musculus* (UCSC mm10), *Equus caballus* (UCSC equCab3), *Bos taurus* (UCSC bosTau9), *Felis catus* (UCSC felCat9), *Canis lupus* (canFam4), *Ericaneus europaeus* (UCSC eriEur2), *Myotis lucifugus* (UCSC Myoluc2), *Loxodonta africana* (UCSC loxAfr3), Dasypus novemcinctus (UCSC dasNov3), Monodelphis domestica (UCSC MonDom5), Aquila chrysaetos (UCSC aquChr2), *Xenopus laevis* (UCSC xenLae2), *Latimeria chalumnae* (UCSC LatCha1), *Callorhinchus milii* (UCSC calMil1), *Petromyzon marinus* (UCSC petMar2), *Petromyzon marinus* (UCSC petMar3), *Eptatretus burgeri* (GenBank GCA_024346535.1), Ciona intestinalis (UCSC ci3).

Additional Primates: *Pan paniscus* (NCBI GCF_029289425.1), *Nomascus leucogenys* (NCBI GCF_006542625.1), *Hylobates moloch* (NCBI GCF_009828535.3), *Colobus angolensis* (NCBI GCF_000951035.1), *Trachypithecus francoisi* (NCBI GCF_009764315.1), *Rhinopithecus roxellana* (NCBI GCF_007565055.1), *Chlorocebus sabaeus* (NCBI GCF_01525025.1), *Theropithecus gelada* (NCBI GCF_003255815.1), *Papio anubis* (NCBI GCF_000264685.3), *Macaca mulatta* (GCF_003339765.1), *Macaca fascicularis* (NCBI GCF_012559485), *Macaca nemestrina* (NCBI GCF_000956065.1), *Cerbocebus atys* (NCBI GCF_000955945.1), Mandrillus leucophaeus (NCBI GCF_000951042.1), Callithrix jacchus (NCBI GCF_011100555.1), *Cebus imitator* (NCBI GCF_001604975.1), *Saimiri boliviensis* (NCBI GCF_016699345.2), *Aotus nancymaae* (UCSC GCF_000952055.2), *Piliocolobus tephrosceles* (NCBI GCF_002776525.5).

Additional Rodents: *Mus spretus* (GenBank GCA_921997135.2), *Mus castaneus* (GenBank GCA_921999005.2), *Mus caroli* (NCBI GCF_900094665.2), *Mus pahari* (NCBI GCF_900095145.1), *Rattus rattus* (NCBI GCF_011064425.1), *Mastomys coucha* (NCBI GCF_008632895.1), *Grammomys surdaster* (NCBI GCF_004785775.1), *Arvicanthis niloticus* (NCBI GCF_011762505.1), *Meriones unguiculatus* (NCBI GCF_002204375.1), *Mesocricetus auratus* (NCBI GCF_017639785.1), *Arvicola amphibious* (NCBI GCF_903992535.2), *Microtus ochrogaster* (NCBI GCF_000317375.1), *Microtus oregoni* (NCBI GCF_018167655.1), *Nannospalax galili* (NCBI GCF_000622305.1), *Jaculus jaculus* (NCBI GCF_020740685.1), *Octodon degus* (NCBI GCF_000260255.1), *Chinchilla lanigera* (NCBI GCF_020740685.1), *Cavia porcellus* (NCBI GCF_000151735.1), *Ictidomys tridecemlineatus* (NCBI GCF_016881025.1).

*Mus musculus* strains (Ferraj et al. 2023): C57/BL6 (GCA_029237415.1), 129S1 (GCA_029237445.1), A/J (GCA_029237185.1), C3H/HeJ (GCA_029237405.1), C3H/HeOuJ (GCA_029237465.1), BALB/cJ (GCA_029237485.1), BALB/cByJ (GCA_029237445.1), NOD (GCA_029234005.1), NZO/HILtJ (GCA_029233705.1), PWD/PhJ (GCA_029234005.1), PWK/PhJ (GCA_029233695.1), WSB/EiJ (GCA_029233295.1), CAST/EiJ (GCA_029237265.1).

### Synteny analysis

For those genomes without assembled *Schlafen* loci, we determined the location of human *PEX12* (NM_000286.3) and *UNC45B* (NM_001267052.2) in contigs or chromosomes. These genes are slowly evolving and could be mapped in most species queried. We then annotated each *Schlafen* homologs relative to: (**1**) their position to the flanking genes; (**2**) other homologs; (**3**) strand orientation. We repeated the same syntenic analysis *SLFNL1* homologs using: *CTPS1* (NM_001905.4) and *SCMH1* (NM_001394311.1).

### Alignment and maximum-likelihood phylogenies

Protein and nucleic acid sequences were aligned using MUSCLE 5.1 (Edgar 2004) or MAFFT 1.5 (Katoh and Standley 2013). We used SMS smart model selection to predict the best model for each alignment (Lefort et al. 2017). Maximum-likelihood phylogenies were built using PhyML 3.3 with either 100 or 500 bootstrap replicates (Guindon et al. 2010). Phylogenetic tree corresponding to SCHLAFEN amino acids sequences in vertebrates was built with JTT model and tree topology optimizing parameters with 500 bootstraps in PhyML. Nucleotides trees was built with the HKY85 model using 100 bootstraps for primate coding sequences and 500 bootstraps for rodent coding sequences.

### Detection of recombination segments

We detected recombination segments in primate and rodent codon alignments using GARD (Kosakovsky Pond et al. 2006) implemented in the HYPHY 2.5.2 package with general discrete model of site to site variation, and three rate classes parameters (Kosakovsky Pond et al. 2020). Recombination segments of <100bp were fused for further analyses.

### Genomic alignments

To compare the genomic sequences of *Slfn12*-related paralogs in *Nannospalax galili* and *Cricetulus griseus* genomes, we extracted ∼16Kb of genomic sequence encompassing the coding loci of *Slfn3* and *Slfn4 or Slfn12_4 and Slfn12_1.* We aligned these sequences using the LAGAN global pair-wise alignment program (Brudno et al. 2003) and visualized them using mVISTA (Frazer et al. 2004).

### Selection analyses

Codon-level dN/dS tests were done using codeml “NSsites”, PAML 4.0 (Yang 2007), with gene trees of degapped Phylip codon alignments. First, we compared NSsites models for evidence of natural selection on full length CDS (“whole gene”, Supplemental Table S2, Supplemental Table S3 and Supplemental Table S4). Statistical significance was measured with χ2 test of twice the difference in log-likelihood between M0 (calculated dN/dS) or its null M0=1 (dN/dS fixed to 1; genetic drift) with 1 degree of freedom. Genes for which we could reject the null (at p-val≤0.05), were tested for positive selection. For this, we used alignments from predicted recombination segments to account for dN/dS variations between gene fragments. In each case, we compared the likelihood of model M8 (allowing for dN/dS>1) with two null models: M7 (not allowing dN/dS>1); and M8a (dN/dS fixed to 1). We used likelihood ratio testing with χ2 to calculate a p-value for positive selection (1 degree of freedom for M8a, 2 for M7). Segments were considered to be subject to positive selection at p-val≤0.05. Sites considered to be under positive selection are those with M8 Bayes Empirical Bayes posterior probability ≥ 99%. Results were consistent between codon frequency 2 or 3 and starting omega of 0,4 or 1,5. All the PAML results presented in Supplemental Table S2 and Supplemental Table S3 are from codon frequency of 2 with starting omega of 0,4. Amino acid coordinates are relative to mouse or human full length CDS orthologs.

We also performed FUBAR analyses on degapped alignments from predicted recombination segments using universal code. Sites considered to be under positive selection are those with a posterior probability ≥ 0.95. (Murrell et al. 2013; Weaver et al. 2018).

### McDonald-Kreitman testing

McDonald-Kreitman test (http://mkt.uab.es/mkt/MKT.asp) on full length degapped CDS alignments from *Slfn8*, *Slfn9* and *Slfn10* using *Mus spretus* as an outgroup for *Slfn9* and *Mus caroli* as outgroup for *Slfn8* and *Slfn10*.

### Species phylogenies

Species divergence time and trees were calculated using timetree.org (Kumar et al. 2017). Species trees were also built using timetree.org specifying a group of species in a list. For more detailed rodent and primate species trees we used previously published phylogenies (Perelman et al. 2011; Steppan and Schenk 2017; Swanson et al. 2019). Mouse strains phylogeny/pedigree was inferred from recent haplotype mapping (Kirby et al. 2010).

### Protein domain annotations

Domain corresponding to (1) the Schlafen core domain (AlbA and Slfn-box); (2) the GTPase-like linker domain (SWADL); and (3) the helicase domain (P-loop NTPase and Walker A and B) were annotated on Human and Mouse proteins according to published structures and predictions (Chen and Kuhn 2019; Garvie et al. 2021; Jo and Pommier 2022; Metzner et al. 2022a; Metzner et al. 2022b).

### Protein structure prediction

Structure predictions of SLFN1, SLFN3 and SLFN4 were calculated with the ColabFold v1.5.2 package which uses AlphaFold2 (Jumper et al. 2021; Mirdita et al. 2022). We used SCHLAFEN protein sequences as queries with default parameters, processed with mmseq2 alignment with the model type set on auto (Steinegger and Soding 2017). To predict homodimer, we used “:” to specify inter-protein chain breaks for modelling dimers and the model type set on alphafold2_multimer_v3. For each protein, we selected the best ranking model based on the structure alignment confidence and evaluated interface confidence using “predicted aligned error” (PAE) measured in Angströms (Å). We rejected dimer predictions for SLFN1 since all the residues had PAE value > 20Å. Conversely, we retained the prediction of SLFN3 and SLFN4 homodimers because most chain A and B residues were < 5Å.

### Structure analysis

We used published pdb (protein data bank format) for human SLFN5 (Metzner et al. 2022a), SLFN11 (Metzner et al. 2022b) and SLFN12 (Garvie et al. 2021) or predicted pdb for rodents. All structures were visualized using ChimeraX (Pettersen et al. 2021). For SLFN5, high sequence similarity allowed us to map rodent positively selected sites on the human structure. To identify of protein interfaces, we used the ChimeraX “contact” option with a 4Å threshold between crystalized or predicted dimer chains.

## DATA ACCESS

All data used and reported in this study can be downloaded freely in public repositories listed in the methods.

## ACKNOWLEDGMENTS

We would like to thank Dr. Lucie Etienne, Dr. Deborah Bourc’his and members of the Molaro laboratory for helpful discussions. This work was funded by the Agence Nationale de la Recherche (ANR): ANR-22-CE12-0006-01; the Fondation pour la Recherche Médicale (FRM): AJE201912009932; and the Institute of Genetics, Reproduction and Development (iGReD).

## REFERENCES

1. Altschul SF, Gish W, Miller W, Myers EW, Lipman DJ. 1990. Basic local alignment search tool. J Mol Biol 215: 403–410.

2. Arslan AD, Sassano A, Saleiro D, Lisowski P, Kosciuczuk EM, Fischietti M, Eckerdt F, Fish EN, Platanias LC. 2017. Human SLFN5 is a transcriptional co-repressor of STAT1-mediated interferon responses and promotes the malignant phenotype in glioblastoma. Oncogene 36: 6006–6019.

3. Bartonicek N, Rouet R, Warren J, Loetsch C, Rodriguez GS, Walters S, Lin F, Zahra D, Blackburn J, Hammond JM et al. 2022. The retroelement Lx9 puts a brake on the immune response to virus infection. Nature 608: 757–765.

4. Basson MD, Wang Q, Chaturvedi LS, More S, Vomhof-DeKrey EE, Al-Marsoummi S, Sun K, Kuhn LA, Kovalenko P, Kiupel M. 2018. Schlafen 12 Interaction with SerpinB12 and Deubiquitylases Drives Human Enterocyte Differentiation. Cell Physiol Biochem 48: 1274–1290.

5. Bell TA, de la Casa-Esperon E, Doherty HE, Ideraabdullah F, Kim K, Wang Y, Lange LA, Wilhemsen K, Lange EM, Sapienza C et al. 2006. The paternal gene of the DDK syndrome maps to the Schlafen gene cluster on mouse chromosome 11. Genetics 172: 411–423.

6. Brudno M, Do CB, Cooper GM, Kim MF, Davydov E, Program NCS, Green ED, Sidow A, Batzoglou S. 2003. LAGAN and Multi-LAGAN: efficient tools for large-scale multiple alignment of genomic DNA. Genome Res 13: 721–731.

7. Bustos O, Naik S, Ayers G, Casola C, Perez-Lamigueiro MA, Chippindale PT, Pritham EJ, de la Casa-Esperon E. 2009. Evolution of the Schlafen genes, a gene family associated with embryonic lethality, meiotic drive, immune processes and orthopoxvirus virulence. Gene 447: 1–11.

8. Chen J, Kuhn LA. 2019. Deciphering the three-domain architecture in schlafens and the structures and roles of human schlafen12 and serpinB12 in transcriptional regulation. J Mol Graph Model 90: 59–76.

9. Cohen-Tannoudji M, Vandormael-Pournin S, Le Bras S, Coumailleau F, Babinet C, Baldacci P. 2000. A 2-Mb YAC/BAC-based physical map of the ovum mutant (Om) locus region on mouse chromosome 11. Genomics 68: 273–282.

10. Crespi B, Nosil P. 2013. Conflictual speciation: species formation via genomic conflict. Trends Ecol Evol 28: 48–57.

11. Daugherty MD, Schaller AM, Geballe AP, Malik HS. 2016. Evolution-guided functional analyses reveal diverse antiviral specificities encoded by IFIT1 genes in mammals. Elife 5.

12. Daugherty MD, Zanders SE. 2019. Gene conversion generates evolutionary novelty that fuels genetic conflicts. Curr Opin Genet Dev 58-59: 49–54.

13. Ding J, Wang S, Liu Q, Duan Y, Cheng T, Ye Z, Cui Z, Zhang A, Liu Q, Zhang Z et al. 2023. Schlafen-5 inhibits LINE-1 retrotransposition. iScience 26: 107968.

14. Ding J, Wang S, Wang Z, Chen S, Zhao J, Solomon M, Liu Z, Guo F, Ma L, Wen J et al. 2022. Schlafen 5 suppresses human immunodeficiency virus type 1 transcription by commandeering cellular epigenetic machinery. Nucleic Acids Res 50: 6137–6153.

15. Eaglesham JB, McCarty KL, Kranzusch PJ. 2020. Structures of diverse poxin cGAMP nucleases reveal a widespread role for cGAS-STING evasion in host-pathogen conflict. Elife 9.

16. Edgar RC. 2004. MUSCLE: multiple sequence alignment with high accuracy and high throughput. Nucleic Acids Res 32: 1792–1797.

17. Ferraj A, Audano PA, Balachandran P, Czechanski A, Flores JI, Radecki AA, Mosur V, Gordon DS, Walawalkar IA, Eichler EE et al. 2023. Resolution of structural variation in diverse mouse genomes reveals chromatin remodeling due to transposable elements. Cell Genom 3: 100291.

18. Fischietti M, Arslan AD, Sassano A, Saleiro D, Majchrzak-Kita B, Ebine K, Munshi HG, Fish EN, Platanias LC. 2018. Slfn2 Regulates Type I Interferon Responses by Modulating the NF-kappa B Pathway. Mol Cell Biol 38.

19. Frazer KA, Pachter L, Poliakov A, Rubin EM, Dubchak I. 2004. VISTA: computational tools for comparative genomics. Nucleic Acids Res 32: W273–279.

20. Gardner A, Ubeda F. 2017. The meaning of intragenomic conflict. Nat Ecol Evol 1: 1807–1815.

21. Garvie CW, Wu X, Papanastasiou M, Lee S, Fuller J, Schnitzler GR, Horner SW, Baker A, Zhang T, Mullahoo JP et al. 2021. Structure of PDE3A-SLFN12 complex reveals requirements for activation of SLFN12 RNase. Nat Commun 12: 4375.

22. Geserick P, Kaiser F, Klemm U, Kaufmann SH, Zerrahn J. 2004. Modulation of T cell development and activation by novel members of the Schlafen (slfn) gene family harbouring an RNA helicase-like motif. Int Immunol 16: 1535–1548.

23. Gubser C, Goodbody R, Ecker A, Brady G, O’Neill LAJ, Jacobs N, Smith GL. 2007. Camelpox virus encodes a schlafen-like protein that affects orthopoxvirus virulence. J Gen Virol 88: 1667–1676.

24. Guindon S, Dufayard JF, Lefort V, Anisimova M, Hordijk W, Gascuel O. 2010. New algorithms and methods to estimate maximum-likelihood phylogenies: assessing the performance of PhyML 3.0. Syst Biol 59: 307–321.

25. Hedges SB, Marin J, Suleski M, Paymer M, Kumar S. 2015. Tree of life reveals clock-like speciation and diversification. Mol Biol Evol 32: 835–845.

26. Hernaez B, Alonso G, Georgana I, El-Jesr M, Martin R, Shair KHY, Fischer C, Sauer S, Maluquer de Motes C, Alcami A. 2020. Viral cGAMP nuclease reveals the essential role of DNA sensing in protection against acute lethal virus infection. Sci Adv 6.

27. Hurst GD, Werren JH. 2001. The role of selfish genetic elements in eukaryotic evolution. Nat Rev Genet 2: 597–606.

28. Jo U, Pommier Y. 2022. Structural, molecular, and functional insights into Schlafen proteins. Exp Mol Med 54: 730–738.

29. Jumper J, Evans R, Pritzel A, Green T, Figurnov M, Ronneberger O, Tunyasuvunakool K, Bates R, Zidek A, Potapenko A et al. 2021. Highly accurate protein structure prediction with AlphaFold. Nature 596: 583–589.

30. Katoh K, Standley DM. 2013. MAFFT multiple sequence alignment software version 7: improvements in performance and usability. Mol Biol Evol 30: 772–780.

31. Kim ET, Dybas JM, Kulej K, Reyes ED, Price AM, Akhtar LN, Orr A, Garcia BA, Boutell C, Weitzman MD. 2021. Comparative proteomics identifies Schlafen 5 (SLFN5) as a herpes simplex virus restriction factor that suppresses viral transcription. Nat Microbiol 6: 234–245.

32. Kim ET, Weitzman MD. 2022. Schlafens Can Put Viruses to Sleep. Viruses 14.

33. Kirby A, Kang HM, Wade CM, Cotsapas C, Kostem E, Han B, Furlotte N, Kang EY, Rivas M, Bogue MA et al. 2010. Fine mapping in 94 inbred mouse strains using a high-density haplotype resource. Genetics 185: 1081–1095.

34. Kosakovsky Pond SL, Poon AFY, Velazquez R, Weaver S, Hepler NL, Murrell B, Shank SD, Magalis BR, Bouvier D, Nekrutenko A et al. 2020. HyPhy 2.5-A Customizable Platform for Evolutionary Hypothesis Testing Using Phylogenies. Mol Biol Evol 37: 295–299.

35. Kosakovsky Pond SL, Posada D, Gravenor MB, Woelk CH, Frost SD. 2006. GARD: a genetic algorithm for recombination detection. Bioinformatics 22: 3096–3098.

36. Kumar S, Stecher G, Suleski M, Hedges SB. 2017. TimeTree: A Resource for Timelines, Timetrees, and Divergence Times. Mol Biol Evol 34: 1812–1819.

37. Kuzmin E, Taylor JS, Boone C. 2022. Retention of duplicated genes in evolution. Trends Genet 38: 59–72.

38. Lefort V, Longueville JE, Gascuel O. 2017. SMS: Smart Model Selection in PhyML. Mol Biol Evol 34: 2422–2424.

39. Li M, Kao E, Gao X, Sandig H, Limmer K, Pavon-Eternod M, Jones TE, Landry S, Pan T, Weitzman MD et al. 2012. Codon-usage-based inhibition of HIV protein synthesis by human schlafen 11. Nature 491: 125–128.

40. Lilue J, Doran AG, Fiddes IT, Abrudan M, Armstrong J, Bennett R, Chow W, Collins J, Collins S, Czechanski A et al. 2018. Sixteen diverse laboratory mouse reference genomes define strain-specific haplotypes and novel functional loci. Nat Genet 50: 1574–1583.

41. Lin YZ, Sun LK, Zhu DT, Hu Z, Wang XF, Du C, Wang YH, Wang XJ, Zhou JH. 2016. Equine schlafen 11 restricts the production of equine infectious anemia virus via a codon usage-dependent mechanism. Virology 495: 112–121.

42. McLaughlin RN, Jr., Malik HS. 2017. Genetic conflicts: the usual suspects and beyond. J Exp Biol 220: 6–17.

43. Metzner FJ, Huber E, Hopfner KP, Lammens K. 2022a. Structural and biochemical characterization of human Schlafen 5. Nucleic Acids Res 50: 1147–1161.

44. Metzner FJ, Wenzl SJ, Kugler M, Krebs S, Hopfner KP, Lammens K. 2022b. Mechanistic understanding of human SLFN11. Nat Commun 13: 5464.

45. Mirdita M, Schutze K, Moriwaki Y, Heo L, Ovchinnikov S, Steinegger M. 2022. ColabFold: making protein folding accessible to all. Nat Methods 19: 679–682.

46. Mitchell PS, Young JM, Emerman M, Malik HS. 2015. Evolutionary Analyses Suggest a Function of MxB Immunity Proteins Beyond Lentivirus Restriction. PLoS Pathog 11: e1005304.

47. Molaro A, Malik HS, Bourc’his D. 2020. Dynamic Evolution of De Novo DNA Methyltransferases in Rodent and Primate Genomes. Mol Biol Evol 37: 1882–1892.

48. Murai J, Tang SW, Leo E, Baechler SA, Redon CE, Zhang H, Al Abo M, Rajapakse VN, Nakamura E, Jenkins LMM et al. 2018. SLFN11 Blocks Stressed Replication Forks Independently of ATR. Mol Cell 69: 371–384 e376.

49. Murrell B, Moola S, Mabona A, Weighill T, Sheward D, Kosakovsky Pond SL, Scheffler K. 2013. FUBAR: a fast, unconstrained bayesian approximation for inferring selection. Mol Biol Evol 30: 1196–1205.

50. Nightingale K, Potts M, Hunter LM, Fielding CA, Zerbe CM, Fletcher-Etherington A, Nobre L, Wang ECY, Strang BL, Houghton JW et al. 2022. Human cytomegalovirus protein RL1 degrades the antiviral factor SLFN11 via recruitment of the CRL4 E3 ubiquitin ligase complex. Proc Natl Acad Sci U S A 119.

51. Perelman P, Johnson WE, Roos C, Seuanez HN, Horvath JE, Moreira MA, Kessing B, Pontius J, Roelke M, Rumpler Y et al. 2011. A molecular phylogeny of living primates. PLoS Genet 7: e1001342.

52. Pettersen EF, Goddard TD, Huang CC, Meng EC, Couch GS, Croll TI, Morris JH, Ferrin TE. 2021. UCSF ChimeraX: Structure visualization for researchers, educators, and developers. Protein Sci 30: 70–82.

53. Pisareva VP, Muslimov IA, Tcherepanov A, Pisarev AV. 2015. Characterization of Novel Ribosome-Associated Endoribonuclease SLFN14 from Rabbit Reticulocytes. Biochemistry 54: 3286–3301.

54. Podvalnaya N, Bronkhorst AW, Lichtenberger R, Hellmann S, Nischwitz E, Falk T, Karaulanov E, Butter F, Falk S, Ketting RF. 2023. piRNA processing by a trimeric Schlafen-domain nuclease. Nature 622: 402–409.

55. Schwarz DA, Katayama CD, Hedrick SM. 1998. Schlafen, a new family of growth regulatory genes that affect thymocyte development. Immunity 9: 657–668.

56. Seong RK, Seo SW, Kim JA, Fletcher SJ, Morgan NV, Kumar M, Choi YK, Shin OS. 2017. Schlafen 14 (SLFN14) is a novel antiviral factor involved in the control of viral replication. Immunobiology 222: 979–988.

57. Stabell AC, Hawkins J, Li M, Gao X, David M, Press WH, Sawyer SL. 2016. Non-human Primate Schlafen11 Inhibits Production of Both Host and Viral Proteins. PLoS Pathog 12: e1006066.

58. Steinegger M, Soding J. 2017. MMseqs2 enables sensitive protein sequence searching for the analysis of massive data sets. Nat Biotechnol 35: 1026–1028.

59. Steppan SJ, Schenk JJ. 2017. Muroid rodent phylogenetics: 900-species tree reveals increasing diversification rates. PLoS One 12: e0183070.

60. Swanson MT, Oliveros CH, Esselstyn JA. 2019. A phylogenomic rodent tree reveals the repeated evolution of masseter architectures. Proc Biol Sci 286: 20190672.

61. Weaver S, Shank SD, Spielman SJ, Li M, Muse SV, Kosakovsky Pond SL. 2018. Datamonkey 2.0: A Modern Web Application for Characterizing Selective and Other Evolutionary Processes. Mol Biol Evol 35: 773–777.

62. Yang JY, Deng XY, Li YS, Ma XC, Feng JX, Yu B, Chen Y, Luo YL, Wang X, Chen ML et al. 2018. Structure of Schlafen13 reveals a new class of tRNA/rRNA-targeting RNase engaged in translational control. Nat Commun 9: 1165.

63. Yang Z. 2007. PAML 4: phylogenetic analysis by maximum likelihood. Mol Biol Evol 24: 1586–1591.

64. Yue T, Zhan X, Zhang D, Jain R, Wang KW, Choi JH, Misawa T, Su L, Quan J, Hildebrand S et al. 2021. SLFN2 protection of tRNAs from stress-induced cleavage is essential for T cell-mediated immunity. Science 372.

65. Zoppoli G, Regairaz M, Leo E, Reinhold WC, Varma S, Ballestrero A, Doroshow JH, Pommier Y. 2012. Putative DNA/RNA helicase Schlafen-11 (SLFN11) sensitizes cancer cells to DNA-damaging agents. Proc Natl Acad Sci U S A 109: 15030–15035.

